# Negatively charged peptide nanofibrils from immunoglobulin light chain sequester viral particles but lack cell-binding and viral transduction-enhancing properties

**DOI:** 10.1101/2020.12.17.423235

**Authors:** Desiree Schütz, Clarissa Read, Rüdiger Groß, Annika Röcker, Sascha Rode, Karthikeyan Annamalai, Marcus Fändrich, Jan Münch

**Affiliations:** Institute of Molecular Virology, Ulm University Medical Center, Ulm, Germany; Central Facility for Electron Microscopy, Ulm University, Ulm, Germany; Institute of Virology, Ulm University Medical Center, Ulm, Germany; Institute of Protein Biochemistry, Ulm University, Ulm, Germany

**Keywords:** peptide nanofibrils, amyloid, infectivity enhancement, fibril-cell interaction

## Abstract

Positively charged naturally occurring or engineered peptide nanofibrils (PNF) are effective enhancers of lentiviral and retroviral transduction, an often rate limiting step in gene transfer and gene therapy approaches. These polycationic PNF are thought to bridge the electrostatic repulsions between negatively charged membranes of virions and cells, thereby enhancing virion attachment to and infection of target cells. Here, we analyzed PNF which are formed by the peptide AL1, which represents a fragment of an immunoglobulin light chain that causes systemic AL amyloidosis. We found that negatively charged AL1 PNF interact with viral particles to a comparable extent as positively charged PNF. However, AL1 PNF lacked cell binding activity and consequently did not enhance retroviral infection. These findings show that virion capture and cell binding of PNF are mediated by different mechanisms, offering avenues for the design of advanced PNF with selective functions.

## Introduction

Self-assembling peptide nanofibrils (PNF) are versatile tools for a growing number of applications in biomedicine^1,2^. Among the most advanced PNF are those that facilitate retroviral gene transfer^3,4^ and two of them, Protransduzin^®3^ and Vectofusin-1^®5^, are commercially available viral transduction enhancers^6^. Protransduzin^®^ fibrils are formed by a 12-mer peptide termed EF-C which is derived from the HIV-1 glycoprotein^3^. EF-C derived PNF (EF-C PNF) are characterized by a high intermolecular ß-sheet content, with lengths between ~100 and 400 nanometer and diameters of 3.4 ± 1.1 nanometer^3^. The EF-C peptide carries two positively charged amino-acid residues (pI: 9.31) and instantaneously forms fibrils upon dilution in polar solvents. The fibrils display a strongly positive zeta potential, indicating a positive surface charge^3^. Vectofusin-1 is a 26-mer cell-penetrating peptide derived from the LAH4 family and forms α-helical PNF structures with sizes of 3 to 10 nanometer^7^, and similar to the EF-C peptide, Vectofusin-1 is positively charged (+4; pI: 10,48). EF-C PNF and Vectofusin-1 enhance retroviral transduction by the same mechanism that has been discovered in studies with naturally occurring fibril-forming peptides in semen. These peptides are proteolytic fragments of the abundant semen proteins prostatic acid phosphatase (PAP) and semenogelins (SEM) 1 & 2, and self-assemble in semen into typical amyloid fibrils with a high level of ß-sheet structure ^3,8–11^. Originally, these fibrils were identified as powerful enhancer of HIV-1 infection which might be exploited by the AIDS-causing virus to enhance its transmission potential during sexual intercourse^8–14^. All semen fibrils are derived from peptides with high isoelectric points, and only the fibrillar but not the monomeric peptides enhance viral infection^8,10,12^. Like EF-C and Vectofusin-1 PNF, PAP and SEM fibrils are polycationic at neutral pH^9^, which enables the fibrils to bind to the negatively charged membranes of both HIV virions and cells, which leads to increased viral attachment to and fusion with target cells^11^. Abrogating the cationic properties of the fibrils through incubation with anionic polymers like heparin or site-directed mutagenesis significantly diminished their ability to enhance infection^11^. Thus, naturally occurring PAP and SEM fibrils as well as engineered EF-C and Vectofusin-1 PNF presumably enhance viral infection/transduction by forming an electrostatic bridge that overcomes the negative repulsions between the viral and cellular membrane, which is normally a rate-limiting step in the viral attachment process^11^. The natural function of semen amyloids is also based on electrostatic interactions: SEVI and SEM fibrils sequester negatively charged apoptotic sperm^15^ and bacterial pathogens^10^ to promote their phagocytotic clearance by macrophages.

To identify other types of amyloid fibrils in the human body that play roles in viral infection or innate immune defense, or may alternatively be applied as transduction enhancer, we here analyzed the 12 mer AL1 peptide derived from the immunoglobulin light chain, which forms polymorphic amyloid-like fibrils *in vitro* ^16–18^ and is of substantial clinical relevance. AL amyloidosis is the most common form of systemic amyloidosis in the Western hemisphere^19^ and caused by extracellular deposition of amyloid fibrils derived from immunoglobulin light chains^20^. The AL1 peptide has a negative pI of 5.52 and does not contain any charged residues. We here show that the amyloid fibrils from AL1 peptide, termed AL1 PNF herein, indeed have a net negative zeta potential and lack cell binding and hence infection/transduction enhancing activity. However, despite the net negative surface charge, AL1 PNF were still capable of sequestering viral particles, suggesting that the overall surface charge of fibrils and virions are not the sole driving force for the formation of fibril-virus complexes. This finding might pave the way for the development of PNF that selectively bind and inactivate enveloped viruses (but not cellular membranes) or that can be tailored with specific cell targeting functionalities, such as antibodies, to allow viral transduction of specific cell types.

## Experimental Section (Materials and Methods)

### Reagents and fibrils

Synthetic peptides PAP248-286 and EF-C were purchased from Celtek peptides (Franklin, Tennessee (TN), USA) or Synpeptide Co., Ltd (Shanghai, China). For fibril formation, peptides were reconstituted and assembled as previously described^3,8,12^. AL1-peptide was chemically synthesized at IZKF Leipzig, Core Unit Peptid-Technologien with the sequence IGSNVVTWYQQL. To prepare fibrils, 5 mg/mL concentration of lyophilized AL1-peptide were dissolved in 50 mM Tris-HCl buffer pH 8.0 and incubated at room temperature for at least 3 days^17^.

### Electron microscopy

Specimens were prepared by placing 5 μl of AL1-PNF suspension onto a formvar and carbon coated 200 mesh copper grid (Plano) followed by a 1 min incubation period at room temperature. The grid was washed three times with water, stained with 2 % (w/v) uranyl acetate solution and dried. Finally, the dryied grid was examined under a JEM-1400 transmission electron microscope (JEOL, Tokyo, Japan) at 120 kV. For cryo-transmission electron microscopy (cryo-TEM) and scanning electron microscopy, AL1-PNF and, for safety reasons, non-infectious env-deleted HIV-1 particles or H_2_O were incubated for 10 min at room temperature prior to adhesion of 3.5 μl to a freshly glow-discharged holey carbon grid. The grids for SEM were blotted, fixed in 1% glutaraldehyde and washed with 5% ethanol in H_2_O. Both, the grids for cryo-TEM and SEM, were then vitrified in liquid ethane by a Vitrobot FP 5350/60 (FEI, Eindhoven, Netherlands). The cryo-TEM grids were analyzed in a JEM – 2100F (JEOL) at 200 kV. For SEM, samples were freeze-dried and platinum (2 nm)/carbon coated (0.5 nm) by electron beam evaporation in a Baf 300 (BAL-TEC AG, Balzers, Principality of Liechtenstein) as described previously (Walther et al., 2008). Grids were analyzed with a Hitachi S-5200 field emission scanning electron microscope (Tokyo, Japan) detecting the secondary electron signal at 10kV.

### Thioflavin T fluorescence

Fibril samples (5-10 μl of 5 mg/ml) were stained with 140-145 μl 50 μM thioflavin T (ThT, Sigma Aldrich Chemie GmbH, Steinheim, Germany) in black 96 well plates with clear bottom (Corning^®^). Fluorescence spectra were recorded on the Tecan Infinite M1000 PRO plate reader (Zürich, Switzerland). 440 nm was chosen as excitation wavelength, emission was detected at 460-600 nm.

### Zeta potential

Surface charge of fibrils/peptides was assessed by mixing 50 μl of a 1 mg/ml solution of fibrils with 950 μl 1 mM KCl solution and measuring in a DTS1061 capillary cell (Malvern, Herrenberg, Germany) using the Zeta Nanosizer and the DLS Nano software Malvern, Herrenberg, Germany). Per sample, three measurements were performed. For neutralization experiments, 0.5 mg/ml AL fibrils were incubated with the indicated substances (50 or 500 μg/ml Heparin/Polybrene), centrifuged for 10 min at 14,000 rpm and resuspended with 50 μl 1 mM KCl. PBS was used as a particle-free control.

### Cell culture and virus generation

Adherent TZM-bl reporter cells (NIH Aids Reagent: TZM-bl from Dr. John C. Kappes, Dr. Xiaoyun Wu and Tranzyme Inc.^21–25^) containing a lacZ reporter gene under the control of a HIV long terminal repeat (LTR) promotor were cultured in DMEM medium supplemented with 120 μg/ml penicillin, 120 μg/ml streptomycin, 350 μg/ml glutamine and 10% inactivated fetal calf serum (FCS) (Gibco, Life Technologies, Frederick, MD). Virus stocks of infectious CCR5-tropic HIV-1 NL4-3 92TH014 were generated by transient transfection of 293T cells with respective proviral plasmids as described (Münch et al., 2007). After transfection and overnight incubation, the transfection mixture was replaced with 2 ml cell culture medium with 2% inactivated FCS. After 40 hr, the culture supernatant was collected and centrifuged for 3 min at 330×g to remove cell debris. Virus stocks were analyzed by p24 antigen ELISA and stored at −80°C. Virus-like-particles of murine leukemia virus tagged with YFP (MLVgag-YFP VLPs) were produced by transfection of HEK293Tcells with pcDNA3_MLV Gag-YFP using Transit LT-1 (Mirus). 2 days post-transfection, supernatants were harvested, clarified by centrifugation, aliquoted and frozen at −80°C until use.

### Cell viability assay

The effect of different fibril concentrations on the metabolic activity of TZM-bl cells was assessed using the ATP-dependent CellTiter-Glo assay (Promega, Madison, Wisconsin (WI), USA). After 3 days incubation with the fibrils, supernatant was discarded and 50 μl PBS and 50 μl CellTiter-Glo Reagent were added to the cells. After incubation and gentle shaking for 10 min at room temperature, luminescence in the cell-free supernatant was determined via the Orion microplate luminometer (Berthold, Bad Wildbad, Germany).

### HIV enhancement assay

To assess the HIV-1 enhancing effect of the different amyloid fibrils, 10^4^ TZM-bl cells in 180 μl cell culture medium were seeded in 96-well flat-bottom plates one day before infection. Concentration series of the different fibrils (0-200 μg/ml) were prepared and then mixed with CCR5-tropic HIV-1 NL4-3 92TH014 (1 ng/ml p24 antigen). After 10 min at room temperature, 20 μl of these mixtures were added to TZM-bl cells and infection rates were determined 3 days post infection by detecting β-galactosidase activity in cellular lysates using the Tropix Gal-Screen kit (Applied Biosystems, Life Technologies, Frederick, MD) and the Orion microplate luminometer (Berthold, Bad Wildbad, Germany). All values represent reporter gene activities (relative light units per second; RLU/s) derived from triplicate infections minus background activities derived from uninfected cells.

### Confocal microscopy

Fibrils (0.5 mg ml^-1^) were stained with Proteostat dye (Enzo Life Sciences, Farmingdale, NY), according to the manufacturer’s instructions (Shen D. et al.,2011). Then, MLV-Gag-YFP particles were added in a ratio of 1:2, resulting in a final fibril concentration of 0.17 mg ml^-1^. After incubating the mixture for 5 min, confocal microscopy was carried out on a Zeiss LSM-710 (Oberkochen, Germany). For analysis of fibril-cell interactions, 20,000 TZM-bl cells were seeded the day before into 1 μ 8 well IBIDI-slides. They were exposed to 5 μg/ml Proteostat-labelled fibrils of the different species (AL1 PNF, SEVI or EF-C PNF) and Hoechst. After incubation for 1 h at 37°C, samples were analyzed before removing the supernatants. Then the supernatants were discarded, cells were washed 3x with 200 μl PBS before 200 μl DMEM without phenol red were added and confocal microscopy was carried out again.

### Flow cytometry

Binding of fluorescent virions to different fibrils was quantified by incubating 250 μl fibrils (final concentration 0.1 mg/ml) with 250 μl of a MLV-Gag-YFP particle suspension with increasing virus titers for 10 min before flow cytometry (BD Canto II, Heidelberg, Germany) was carried out. To exclude that the increase in fluorescence is due to unspecific effects, different concentrations of MLV-Gag virions lacking the YFP expression cassette were tested for fibril binding in parallel, using a constant AL fibril concentration of 0.1 mg/ml. For analyzing cell binding of the fibril species over time, Proteostat stained SEVI, EF-C PNF and AL1 PNF fibrils were added to a final concentration of 5 μg/ml to 200,000 TZM-bl cells seeded the day before into 12-well plates. After certain incubation times, supernatants of the cells were discarded, cells were washed 3x with PBS, trypsinated, transferred to FACS tubes, centrifuged and washed with PBS again. Then, cells were fixed with 500 μl 2% PFA and analyzed for bound fibrils via the red fluorescence channel using flow cytometry. For normalization, the complete fluorescence intensity of fibrils was divided by complete fluorescence of DAPI to get fibrils / cell.

### p24 ELISA

To quantify the interaction of virions with target cells, 5,000 TZM-bl cells were seeded one day before into 96-well plates in a volume of 50 μl. The next day, 40 μl of different fibril concentrations were incubated with 40 μl HIV-1 dilution (40 ng/ml p24 antigen) for 10 min and 20 μl of these mixtures were added to the cells in triplicates. After 3 h incubation at 37°C, unbound virus was removed and cells were washed 3x with DMEM before they were lysed with 1% Triton X-100 for 1 h at 37’C. Cell associated HIV-1 capsid antigen was detected using an in-house p24 ELISA.

## Results and Discussion

### AL1 peptide forms negatively charged PNF lacking infection enhancing activity

AL1 PNF were analyzed by TEM confirming the formation of twisted fibrils with diameters varying roughly between 8.0 and 22.0 nm (Figure 1A), due to fibril polymorphism^18^. Amyloid formation was further corroborated by changes in the fluorescence intensities of the amyloid-specific dye Thioflavin T (Figure 1B). To further characterize and compare the fibrils to the seminal amyloid PAP248-286, we performed zeta potential measurements of both native peptides and fibrils. AL1 PNF have an isoelectric point of 5.52 and featured a net negative surface charge (−12 mV) whereas PAP248-286 fibrils were positively charged (16 mV) in KCl solution, confirming previous data^3,10,12,26^. Native AL peptide features small negative (−3 mV) and PAP284-286 minor positive (0.3 mV) surface charge (Figure 1C). Due to the rapid assembly of EF-C peptide into fibrils^3^, zeta potentials of freshly dissolved peptide could not be determined, however, the assembled EF-C PNF were positively charged (+ 12 mV), as shown before^3^.

**Figure 1:**
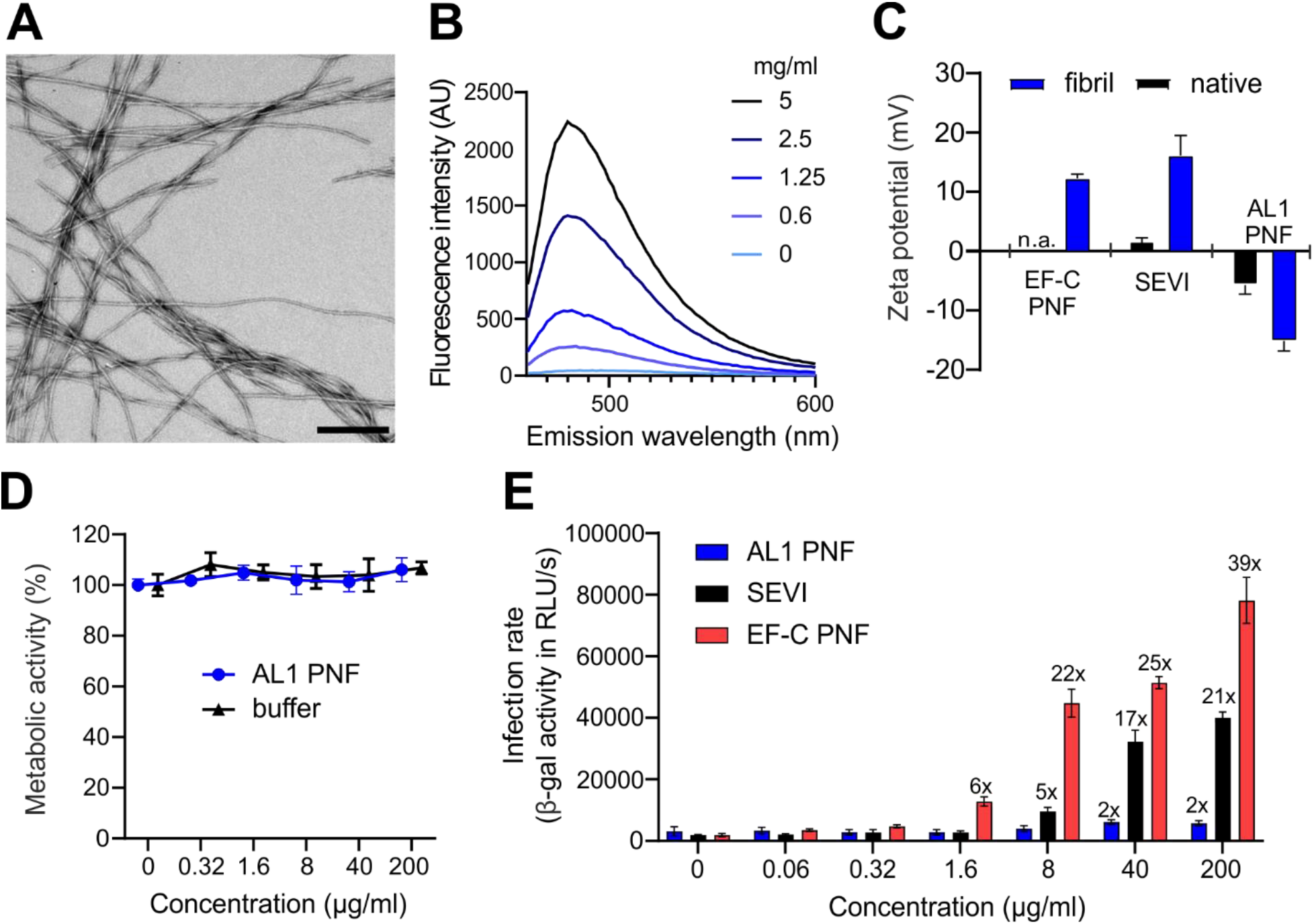
AL1 peptide forms negatively charged fibrils that do not enhance infection. A) Transmission electron microscopy image of AL1 PNF, Scale bar: 250 nm. B) ThT emission spectrum of indicated concentrations of AL1 fibrils. C) Zeta potential of freshly dissolved and fibrillar PAP248-286 and AL1 peptides, and fibrillar EF-C PNF. D) Metabolic activity of TZM-bl cells incubated for three days with indicated concentrations of AL1 PNF or buffer only. Values represent mean values ± SD relative to the untreated control. E) AL1 PNF lack HIV-1 enhancing activity. SEVI, EF-C and AL1 PNF were incubated with HIV-1 and mixtures were added to TZM-bl reporter cells. After 3 days, infection rates were determined. Values represent ß-galactosidase activities obtained from triplicate infections ±SD (RLU/s: relative light units per second).

Before studying the impact of ALI PNF on infection, we verified that they did not impair cellular metabolic activity at concentrations of up to 200 μg/ml (Figure 1D). We then determined the infection enhancing activity of AL1 PNF, and as controls, PAP248-286 fibrils (SEVI) and EF-C PNF. As shown in Figure 1E, EF-C PNF and SEVI fibrils enhanced viral infection in a concentration-dependent manner, with up to 39-fold increased infection rates for EF-C PNF, and 21-fold increased infectivity in the presence of SEVI, at the highest tested concentration. On the contrary, AL1 PNF only marginally increased viral infection (2-fold) at 40 and 200 μg/ml. Thus, AL1 fibrils lack potent infection enhancing activity, presumably due to the negative surface charge which may prevent the interaction with viral and cellular membranes.

### AL1 PNF bind viral particles

To prove that AL1 PNF featuring a negative zeta potential indeed lack virion binding activity, we performed scanning electron microscopy and cryo-transmission electron microscopy of fibrils that were exposed to non-infectious env-deleted HIV-1 or MLV particles. Unexpectedly, our analysis showed that virions were sequestered by the AL1 PNF network (Figure 2A and B). Individual fibrils aligned to the viral envelope bilayer (Figure 2A and B) without causing obvious alterations in the viral membrane (Figure 2B). Confocal microscopy confirmed this finding and revealed complex formation of virions and fibrils (Figure 2 C), very similar to those previously obtained for SEVI^8^, Semenogelin^27^ and EF-C PNF^3^. Virion binding by AL1 PNF, EF-C PNF and SEVI fibrils was further quantified by flow cytometry. For this, a constant concentration of fibrils (100 μg/ml) was incubated with increasing concentrations of the fluorescently labeled retroviral particles, and complex formation was analyzed via flow cytometry. A VLP-dose dependent increase in fluorescence was observed for all fibril species (Figure 2D and E). Interestingly, the fluorescence signals for AL1 fibrils were significantly higher than for the respective EF-C or SEVI-fibril/virion complexes (Figure 2D). However, we found AL1 PNF to exhibit autofluorescence, giving rise to higher signals. In order to exclude unspecific signals from fibrils with increasing virion concentrations, we compared fluorescence signals of AL1 PNF and fluorescent retroviral particles with AL1 PNF incubated with the same concentrations of non-fluorescent virions. For AL1 fibrils and non-fluorescent virions, no fluorescence increase of fibrils with increasing virion concentrations was observed, proving the specificity of signals (Figure S1).

**Figure 2:**
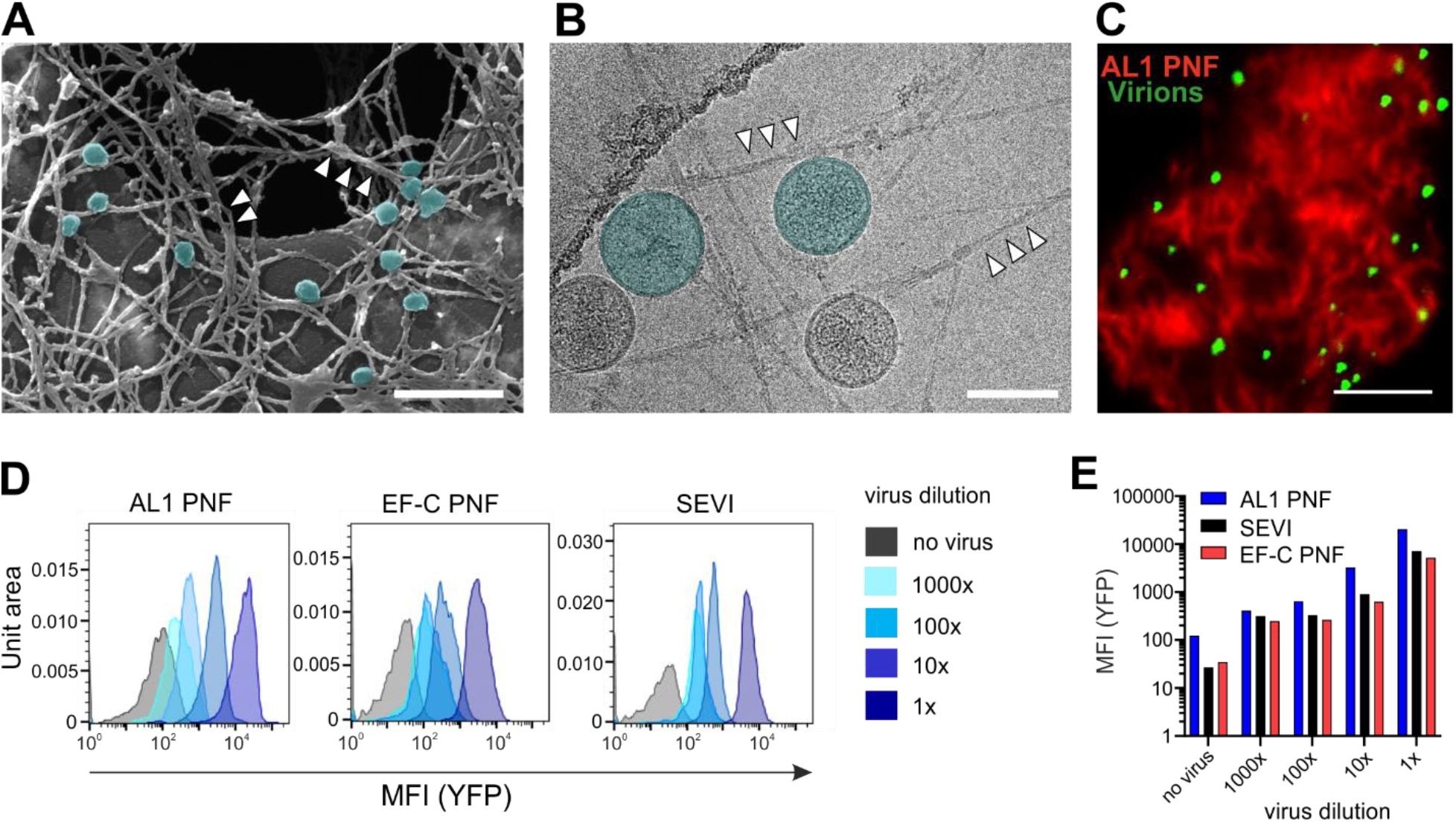
AL1 PNF bind retroviral particles. A: Scanning electron microscopy and B: Cryo-TEM of AL1 PNF (*white arrows*) incubated with retroviral MLV-Gag particles (*blue*; not all particles are colored to see detailed structures). Scale bars: 500 nm (A) and 100 nm (B). C: Confocal microscopy images of AL1 PNF stained with Proteostat dye (*red*) and incubated for 5 min with retroviral MLV-Gag-YFP particles (*green*); scale bar: 5 μm. D: Representative FACS histograms of fibrils incubated with different MLV-Gag YFP dilutions. E: Flow cytometry analysis of fibrils incubated with serial dilutions of fluorescent retroviral particles.

### Low cell binding activity of AL1 PNF as compared to EF-C PNF and SEVI

AL1 PNF bind virions but do not enhance viral infectivity. Thus, we next analyzed if AL1 PNF are capable of binding to the cell membrane, as previously reported for EF-C PNF^3^ and SEVI^8^. To test this, cells were incubated with the three Proteostat-labelled fibril species for 1 h, and then analyzed by confocal microscopy prior to or after thorough washing. As shown in Figure 3A, the three labeled fibril species were readily detectable as red-fluorescent patches on the cell surface. Washing hardly affected EF-C PNF fluorescence, indicating a strong interaction of these fibrils with the cell surface (Figure 3A). Washing removed only some of the SEVI fibrils while AL1 PNF fluorescence disappeared (Figure 3A). Flow cytometry results confirmed these findings: after extensive washing, almost all EF-C PNF and ~ 61% of SEVI remained cell associated. In contrast, more than 80% of the AL1 PNF were removed by washing. We next analyzed whether changes in the incubation time may affect AL1 PNF binding. However, even after 2 hours of incubation, AL1 PNF could be easily removed from the cell surface, whereas SEVI and EF-C fibrils remained bound (Figure 3C). Thus, AL1 PNF do not efficiently attach to the cell surface which may explain the lack of viral infection enhancing activity.

**Figure 3:**
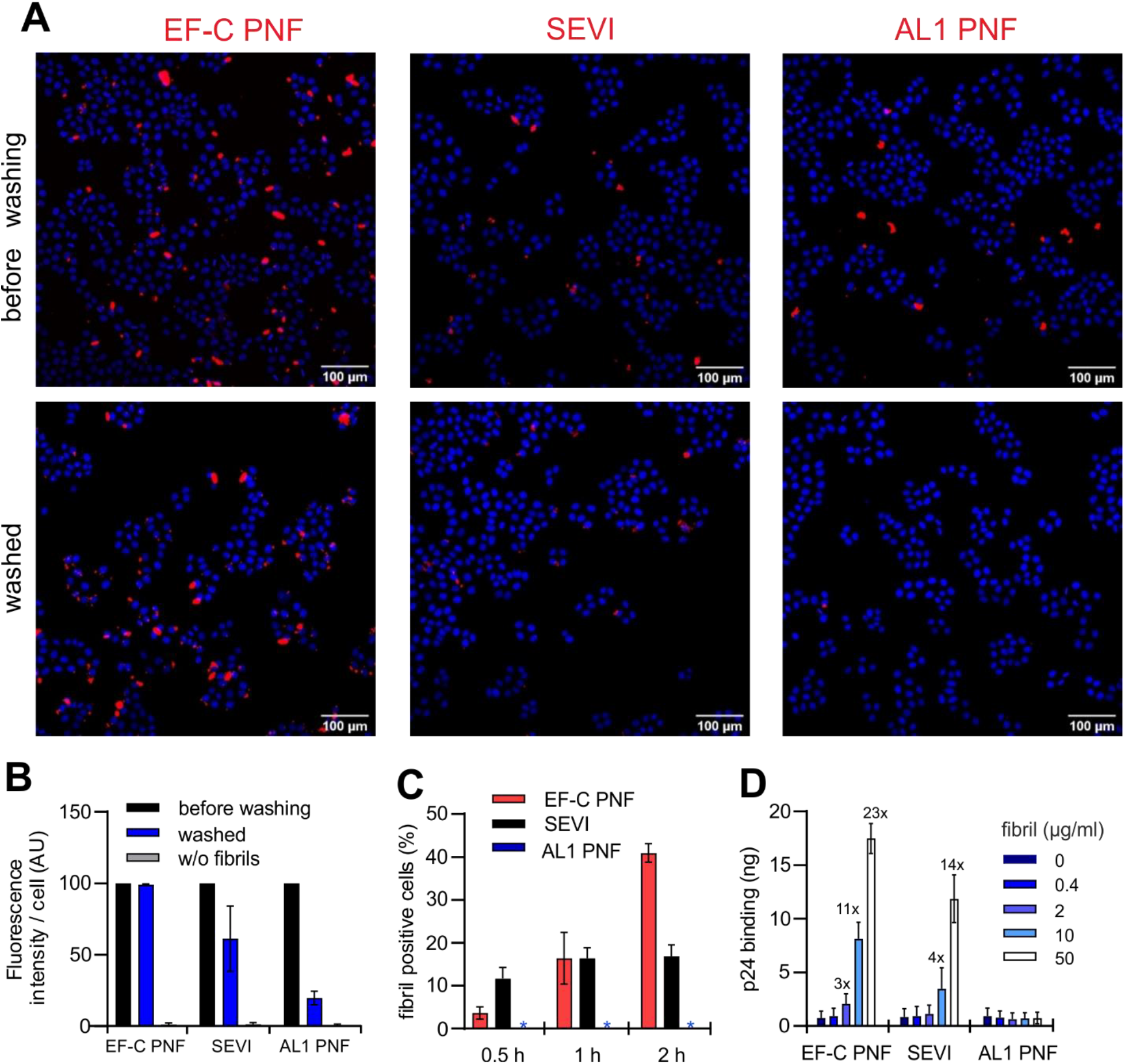
AL1 PNF lack cell binding activity. A: Representative confocal microscopy images of TZM-bl HeLa cells incubated with Proteostat-labeled (red) EF-C, SEVI and AL1 PNF (each 5 μg/ml). After one hour, cells were imaged, washed 3 times in PBS and imaged again. Nuclei were stained with Hoechst (blue). B: Quantitative evaluation of Proteostat-labeled fibril binding using ImageJ from two individual experiments with 3-4 images each using ImageJ. Fluorescence intensity was normalized to intensities obtained before washing. C: Flow cytometry evaluation of Proteostat-labeled fibril binding to TZM-bl HeLa cells. TZM-bl HeLa cells were incubated with the Proteostat-labelled fibrils, washed, trypsinized and analysed by flow cytometry. Shown are means (±SD) of two independent experiments. Asterisks indicate background signal only. D: TZM-bl HeLa cells were inoculated with HIV-1 that was exposed to indicated concentration of the fibrils. After 3 hours, cells were washed 3x times, lysed and analyzed for viral capsid antigen by p24 ELISA. Shown are means (±SD) of two independent experiments.

To corroborate this finding, we quantified the amount of virus that was delivered to the cellular membrane by the three types of fibrils using a p24 antigen ELISA that quantifies the viral capsid protein. As expected ^8^, SEVI and EF-C PNF resulted in a dose-dependent increase of cell-bound viral capsid antigen (Figure 3D). Cell-associated p24 was increased up to 23-fold upon virus incubation with EF-C PNF and up to 14-fold with SEVI. However, no increase in virus binding was observed with AL1 PNF (Figure 3D). Collectively these findings demonstrate that AL1 PNF effectively sequester viral particles but lack cell binding and consequently viral infection enhancing activity.

## Discussion and conclusions

To identify functional amyloids of the human body that may play roles in innate immunity or be developed as nanotechnological tools, we here analyzed amyloid-like fibrils formed by a specific peptide derived from the antibody light chain, the AL1 peptide^17^. In contrast to viral infection enhancing fibrils such as SEVI or EF-C that have a positive surface charge^3,8,10,11^, AL1 PNF feature a net negative surface charge. AL1 PNF lacked viral infection enhancing activity, which was expected because the anionic fibrils might not bridge or neutralize the negative charge repulsions that exist between viral and cellular membranes^6,11^. In fact, no binding to the cellular membrane was observed for AL1 PNF, in contrast to cationic EF-C and SEVI fibrils. It came to a surprise, however, that AL1 PNF featured no deficit in virion binding, and interacted with viral particles to a similar extent as positively charged EF-C and SEVI fibrils. Thus, fibrils with a net negative charge are also capable of sequestering viral particles and the net positive surface charge of fibrils is not the only driving force for binding to virions. However, it needs to be considered that AL1 PNF may feature positively charged surface patches which mediate electrostatic interactions with negatively charged virions. In fact, a recently established model of AL1 PNF suggests the presence of positive charges at the fibril surface (Figure S2). Thus, even though the fibril has a net negative net charge, cationic regions on the fibril surface may be sufficient for virion capture, but not for cell binding.

The selective interaction of AL1 PNF with membranes of virions but not cells suggests that differences in the composition of the viral and cellular lipid bilayer may account for this observation. The retroviral envelope is derived from the cell membrane which resembles a typical lipid bilayer and is negatively charged due to the anionic phospholipid phosphatidylserine^28,29^. However, viral and cellular membranes are not identical. First, embedded into the viral membrane are viral glycoproteins that mediate attachment and infection of target cells and may serve as binding partners of AL1 PNF. However, as shown by TEM, Cryo-EM, confocal microscopy and flow cytometry, AL1-PNF fibrils captured lentiviral or retroviral particles lacking viral glycoproteins, which were used in these experiments for safety reasons as they are not infectious. Second, retroviral particles are budding through membrane microdomains which are rich in cholesterol, sphingolipids, as well as GPI-linked and fatty acylated proteins ^29,30^. Consequently, the viral membrane is enriched in these fatty acids. Notably, a recent article showed that HIV-1 particles are also heavily enriched in phosphoinositides, in particular phosphatidylinositol 4,5-bisphosphate (PIP2) and phosphatidylinositol (3,4,5)-trisphosphate (PIP3)^31^. PIP3 is the most negatively charged plasma membrane lipid^32^ and might render viral particles much more polyanionic than the cytoplasmic membrane. Consequently, the highly negatively charged viral membrane may allow binding into cavities between the positive patches in AL1 PNF, whereas AL1 PNF are not electrostatically attracted by cells with lower charge density, explaining the lack of viral enhancing activity.

Amyloid fibrils have long been associated with diseases but it became increasingly evident that fibrils also exert “functional” roles^8,33,34^. One example of functional amyloid in humans are fibrils formed by fragments of PAP or Semenogelin which are naturally present in semen and are responsible for removal of apoptotic sperm and bacterial pathogens in the female reproductive tract^35,36^. Other examples are amyloidogenic amyloid-β-peptide variants which are involved in Alzheimer’s disease but have recently been shown to induce microbial agglutination and to exert antimicrobial and antiviral activity^37–39^. Even though AL1 PNF sequestered viral particles, the fibrils did not abrogate viral infectivity, even at elevated concentrations. Whether AL1 PNF serve any functional role, e.g. antibacterial activity, needs to addressed in follow up studies.

Our results show that binding of fibrils to virions and cells underlies shared but also separable mechanisms and are seemingly more complex than previously anticipated. However, the identification of fibrils that selectively bind viral particles without enhancing viral infection may open novel avenues for research. For example, AL1 PNF equipped with membrane destroying agents such as molecular tweezers^26,40,41^ may act as direct antiviral nanomaterials that specifically target the viral envelope and not the cell. Alternatively, AL1 PNF carrying specific cell targeting moieties may allow selective delivery of viral vectors or drugs to specific cell types or tissues^34,42^.

## Author Contributions

D.S. and A.R. performed all viral experiments and ThT measurements, C.R. did SEM and cryo EM analyses; R.G. measured zeta potentials; S.R. supported D.S. in performing experiments; K.A. and M.F. did TEM, helped C.R. in cryo EM and generated the AL1 PNF model, J.M. conceptualized and supervised the work and wrote with D.S. the manuscript. All authors have given approval to the final version of the manuscript.

## Acknowledgments

We thank Matthias Schmidt and Irene Vázquez Martín (Ulm University, Institute for Protein Biochemistry) for technical support. This project has received funding through the German Research Foundation (CRC 1279, project A03 to J.M. and M.F. and FOR 2969 to M.F. and C.R.), the Leibniz Foundation (Controlling and Switching of Function of Peptide and Protein based BioSurfaces: From Fundamentals to Applications; project to J.M.) and the Volkswagen Stiftung (to J.M.). Desiree Schütz and Rüdiger Groß are part of the International Graduate School in Molecular Medicine Ulm. Rüdiger Groß acknowledges funding by a scholarship of International Graduate School in Molecular Medicine Ulm.

## Supporting Information

**Figure S1:**
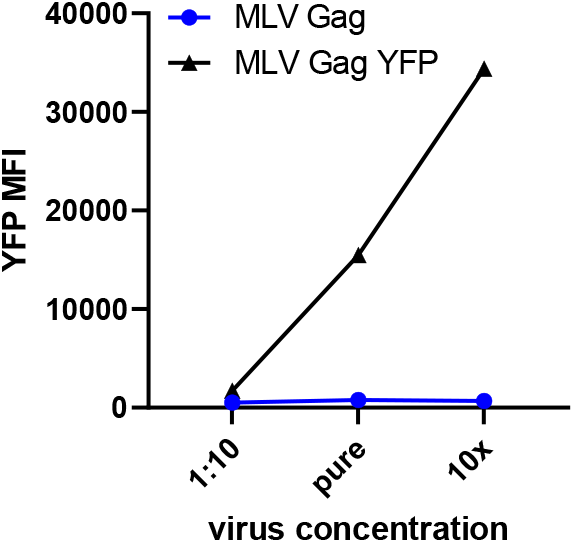
AL1 PNF fluorescence increases upon incubation with increasing concentrations of MLV Gag YFP but not with MLV Gag virions. Equal concentrations of MLV Gag YFP and MLV Gag virions were incubated with 0.1 mg/ml of AL-PNF and analyzed via flow cytometry.

**Figure S2:**
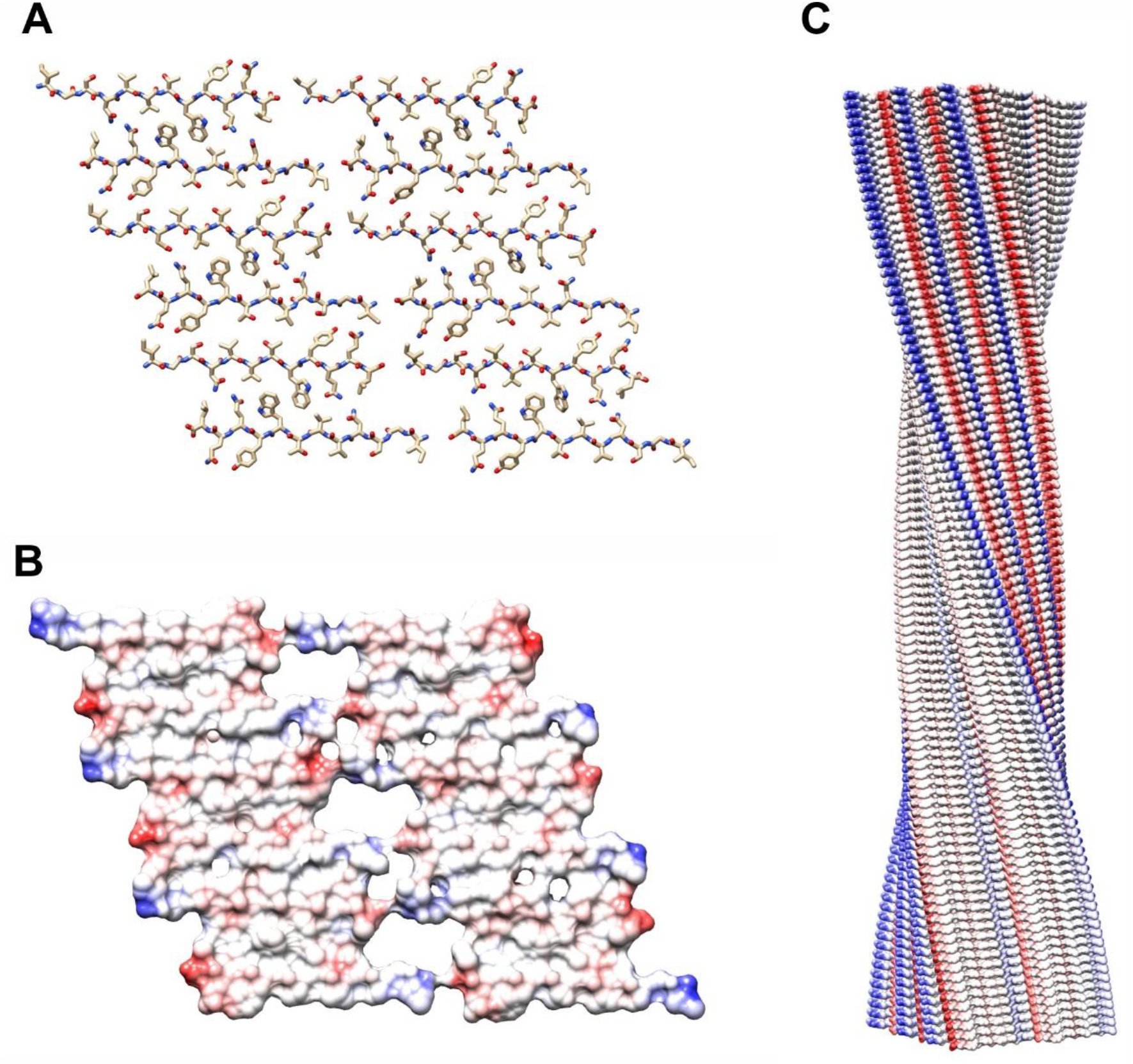
Structural model of the main fibril morphology formed by AL1 peptide. A: Stick representation of a structural model of a cross-sectional layer of the main AL1 PNF morphology as published previously^17^. Red: oxygen; blue: nitrogen; opal: carbon; hydrogens omitted. B: Coulombic surface coloring of the cross-sectional slice. Red: negative charge; blue: positive charge. C: Side view of a fibril segment containing 100 peptide layers shown with Coulombic surface coloring.

